# Robust Normalization of Luciferase Reporter Data

**DOI:** 10.1101/673087

**Authors:** Andrea Repele, Manu

**Author notes:** JMB–Center for Immunity and Immunotherapies, Seattle Children’s Hospital, Seattle, WA, United States of America.

## Abstract

Transient Luciferase reporter assays are widely used in the study of gene regulation and intracellular cell signaling. In order to control for sample-to-sample variation in luminescence arising from variability in transfection efficiency and other sources, an internal control reporter is co-transfected with the experimental reporter. The luminescence of the experimental reporter is normalized against the control by taking the ratio of the two. Here we show that this method of normalization, “ratiometric”, performs poorly when the transfection efficiency is low and leads to biased estimates of relative activity. We propose an alternative methodology based on linear regression that is much better suited for the normalization of reporter data, especially when transfection efficiency is low. We compare the ratiometric method against three regression methods on both simulated and empirical data. Our results suggest that robust errors-in-variables (REIV) regression performs the best in normalizing Luciferase reporter data. We have made the `R` code for Luciferase data normalization using REIV available on GitHub.

## 1. Introduction

Transient reporter assays are an important and widely used tool in the study of gene regulation [1–4], intracellular cell signaling [5–7], and other areas of molecular, cellular, and developmental biology [8–10]. In a reporter assay, the activity of a promoter, potentially in combination with an enhancer, is measured by placing an inert reporter gene such as *luciferase* or *lacZ* under the control of the promoter. *luciferase* is commonly used as the reporter gene since the assay has both high sensitivity and a very high dynamic range [11].

An important technical challenge in the analysis of Luciferase reporter data is that transfection efficiency can vary significantly from sample to sample, especially in hard to transfect cell types such as primary cells. Other sources of random experimental error, such as the amount of transfected DNA, number of cells assayed, cell viability, and pipetting errors further compound transfection efficiency variation to yield Luciferase luminescence data that can vary over an order of magnitude [4,8,12]. For example, a reporter of *CCAAT/Enhancer binding protein, α* (*Cebpa*) promoter expression exhibits ∼3-fold variation in luminescence when assayed in PUER cells [13–15], a myeloid cell line with low transfection efficiency (Fig 1; Section 3.1).

**Figure 1.**
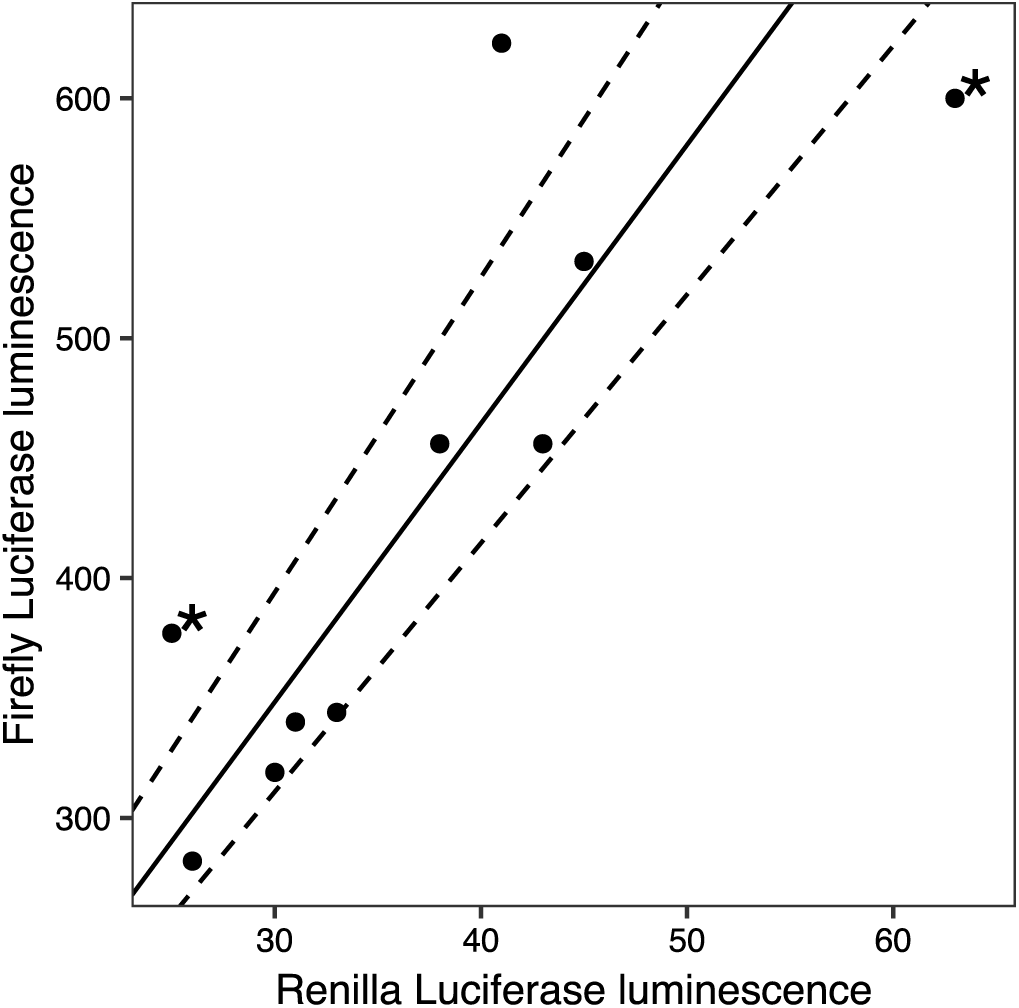
Example firefly and *Renilla* luminescence data from a myeloid cell line. Firefly *luciferase* was under the control of the *Cebpa* promoter, while *Renilla luciferase* was under the control of the CMV promoter. Luminescence is reported in relative luminescence units (RLUs). The best-fit line (solid) determined by robust errors-in-variable (REIV) regression is shown. Dashed lines represent the 95% confidence interval for slope determined by bootstrapping (Section 3.4.4). Potential outliers are indicated with asterisks.

The prevalent method of correcting for sample-to-sample luminescence variation [3,8] is to co-transfect an independent control reporter, such as *Renilla luciferase* expressed from a constitutive promoter, along with the promoter/enhancer reporter being assayed. Since transfection efficiency and other experimental errors would affect both constructs equally, the activity of the experimental promoter relative to the constitutive promoter can be determined by taking the ratio of firefly and *Renilla* luminescence. Let there be *i* = 1, …, *N* samples and *F*_*i*_ and *R*_*i*_ represent firefly and *Renilla* luminescence in the *i*th replicate. Then, the relative activity is estimated as

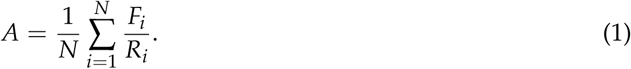

Although it is in common use, “ratiometric” normalization is statistically unsound since it weights low- and high-luminescence replicates equally even though the former produce less reliable estimates of normalized reporter activity. This problem is especially acute in cell types having low transfection efficiency and, consequently, high variability in luminescence measurements.

Here we propose an alternative approach to the normalization of Luciferase reporter data that is more robust than the ratiometric method. Firefly luminescence is expected to be proportional to *Renilla* luminescence,

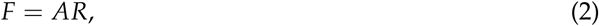

and therefore, the relative activity *A* can be estimated as the slope of the best fit line using linear regression (Fig. 1). Ordinary least-squares (OLS) regression places a higher weight on high-luminescence points and thus avoids the main weakness of ratiometric normalization.

We also considered two alternatives to OLS to find the method best suited for the normalization of Luciferase data. OLS assumes that the values of the independent variable, *Renilla* luminescence in our case, are known exactly and don’t include random errors. Since *Renilla* luminescence is itself a random variable in transient assays, we evaluated errors-in-variables (EIV) regression (Section 3.4.3; [16]). In EIV regression, the loss function is the sum of the squares of the orthogonal distance of each data point from the line, leading to the minimization of errors in both the independent and the dependent variables. Lastly, we considered robust errors-in-variables (REIV) regression (Section 3.4.4; [17]). Luciferase luminescence data contain potential outliers (Fig. 1; [4,8]), which can unduly influence activity estimates. In REIV, the estimation of the slope and intercept is rendered insensitive to outliers by utilizing a bounded loss function.

In this study we compared the four methods described above, ratiometric, OLS, EIV, and REIV, in two ways. First, we applied each method to simulated data and assessed its ability to recover the “true” activity used to generate the synthetic data. Second, we tested the methods on empirical measurements of *Cebpa* promoter activity in PUER cells. Our results show that while ratiometric normalization performs poorly on high variability data from low transfection efficiency experiments, REIV performs the best and robustly estimates activity under a variety of different conditions.

## 2. Results and Discussion

We evaluated the performance of four methods, ratiometric, OLS, EIV, and REIV, on simulated data, where the “ground truth” is known. Reporter assays in cells with low transfection efficiency yield activity data with a high coefficient of variation, making normalization more challenging. Other sources of experimental error impact normalization similarly, and accurate normalization becomes more challenging with increasing standard deviation of the experimental error. We aimed to evaluate the methods’ ability to recover the known promoter activity at different levels of transfection efficiency and experimental error.

We simulated sample-to-sample variation in transfection efficiency (*t*) (Section 3.5) by sampling from the Beta distribution. The coefficient of variation of the Beta distribution is inversely related to mean transfection efficiency [16] and consequently synthetic data from low transfection efficiency simulations have greater variability, mirroring experimental reality. Mean transfection efficiency 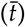 was varied to assess how the methods perform as sample-to-sample variability increases. Random experimental errors, modeled with a Gaussian mixture model [17], were introduced into both *Renilla* and firefly simulated data (Section 3.5). The Gaussian mixture model includes low-frequency contamination by errors with a higher standard deviation and allowed us to simulate outliers (Fig. 1). We varied the standard deviation of *Renilla* errors (*σ*_11_) to assess the performance of each method at different levels of experimental variability. Robustness of each method to outliers was assessed by varying the standard deviation of the contaminating errors (*σ*_12_). We also varied the sample size (*N*) and “true” activity level to investigate how they impact normalization.

Simulated data were generated for each combination of activity, transfection efficiency, sample size, and error standard deviations. Each normalization method was used to estimate activity from the exact same set of simulated data. The simulations were repeated *M* = 300 times and the performance of the methods was assessed using three different measures. Let the known or true activity be *A* and the activity estimated in the *k*th simulation be *A*_*k*_. We utilized the relative absolute difference between the true and estimated values,

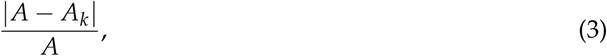

as a measure of the relative bias in the estimation. Second, we computed the relative median absolute deviation (MAD) of the estimated activities *A*_*k*_ as a measure of precision. Although readily interpretable, activity based measures are less reliable when the true activity is high. Since the activity is the slope of the regression line, small deviations of the regression line from the true line lead to large deviations in activity when the true activity is large. For this reason, we also utilized a measure developed by Zamar (1989) [17],

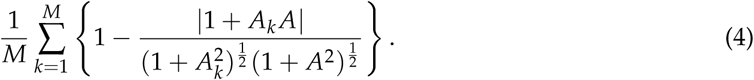

The “Zamar criterion” is independent of the activity level since it is invariant under rotations of the *xy*-plane [17] and serves well to compare different methods. The Zamar criterion takes values between 0 and 1, the former implying perfect recovery of the true activity.

In the first set of simulations, we assessed the ability of the methods to infer the activity from simulated data lacking outliers (*σ*_12_ = *σ*_11_) at different levels of mean transfection efficiency. The performance of all methods improved with increasing transfection efficiency (Fig. 2(a)), however the regression-based methods, OLS, EIV, and REIV, performed better at lower transfection efficiencies as compared to the ratiometric method. At transfection efficiencies less than 25%, the ratiometric method produces rather inaccurate estimates of activity, and the 90th percentile of relative bias is ∼300% at 10% transfection efficiency. Next, we investigated whether the accuracy of the estimates could be improved by increasing sample size (*N*). Since the main difference between the ratiometric and the other methods arises at low transfection efficiencies, we set the mean transfection efficiency to 25% for all subsequent simulations. Whereas the performance of the regression based methods improved with increasing sample size, the bias in the activity inferred by the ratiometric method did not decrease with sample size (Fig. 2(b)). These set of simulations suggested that the ratiometric method of normalization results in biased activity estimates that do not improve even at very large sample sizes.

**Figure 2.**
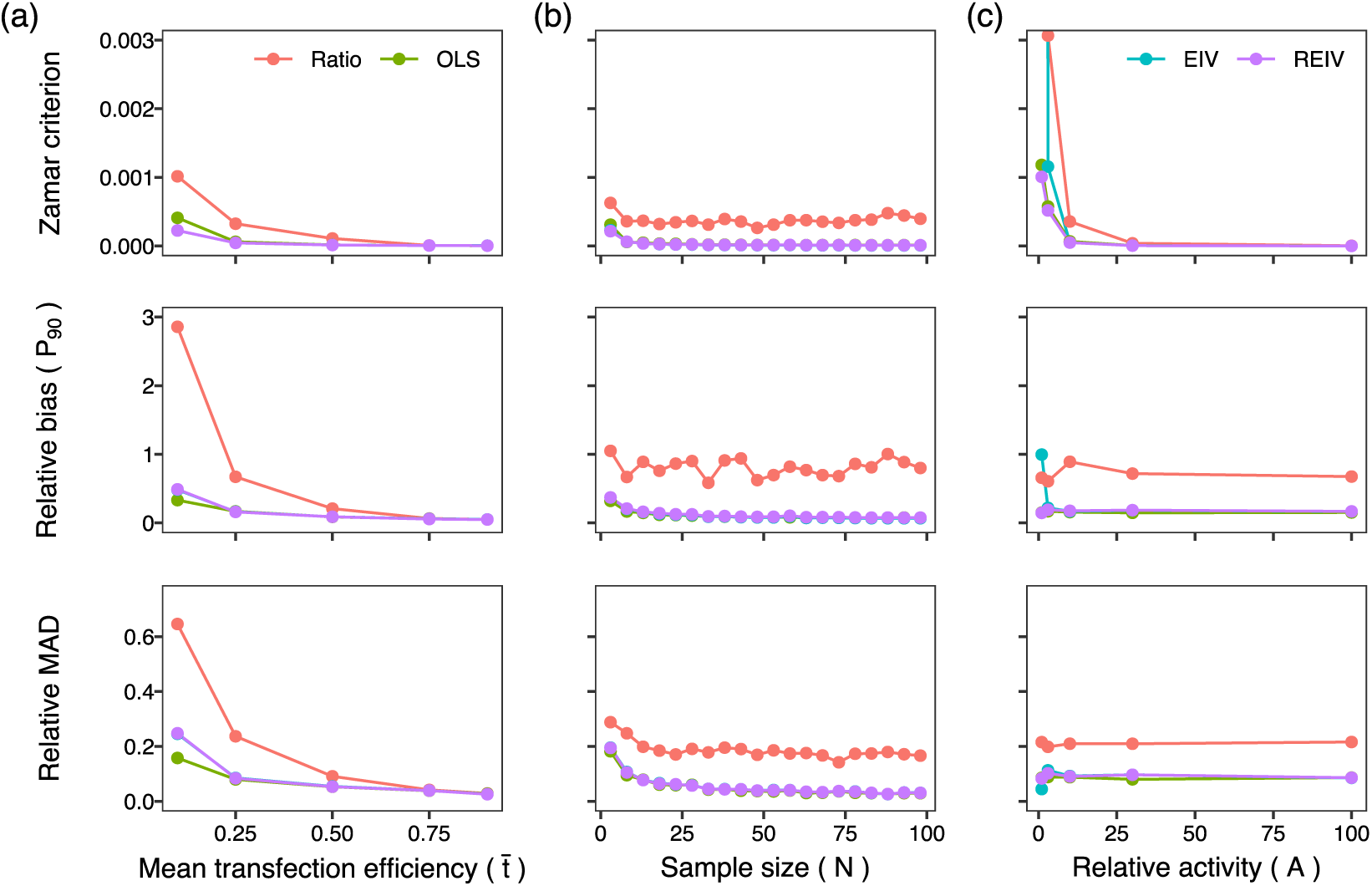
Tests of normalization methods on simulated data. The top panels plot the Zamar criterion, which is 0 for perfect inference of the true activity. The middle panels plot the 90th percentile (*P*_90_) of the relative bias. The bottom panels plot the relative median absolute deviation, a measure of precision. (**a**) Performance of the methods plotted against mean transfection efficiency. True activity: *A* = 10. Sample size: *N* = 10. *Renilla* errors: *σ*_11_ = *σ*_12_ = 3. Firefly errors were scaled with activity: *σ*_21_ = *σ*_22_ = 10*σ*_11_. (**b**) Performance plotted against sample size. Mean transfection efficiency: 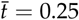. The other parameters are the same as in panel (**a**). (**c**). Performance plotted against true relative activity. Sample size: *N* = 10. The other parameters are the same as in panel (**b**).

In the third set of simulations, we investigated the effect of the activity level on the normalization (Fig. 2(c)). The performance of all the methods deteriorated at activity levels close to 1. An activity level of 1 implies that firefly luminescence is comparable to *Renilla* luminescence, which only occurs for exceptionally weak promoters since it is common practice to transfect the firefly plasmid in a 10- to 200-fold excess over the *Renilla* plasmid. Next we investigated the effect of experimental error in *Renilla* luminescence (Fig. 3(a)). As one would expect, the performance of all methods deteriorates as the magnitude of experimental error is increased. Furthermore, the ratiometric method had the worst performance amongst all the methods. The errors-in-variables methods, EIV and REIV, were the most resilient to errors in *Renilla* (Fig. 3(a); Zamar criterion) since OLS does not attempt to minimize errors in the independent variable. Finally, we investigated the performance of the methods in the presence of outliers. So far, the simulations had been carried out without outliers since the standard deviation of the contaminating errors was the same as the Renilla experimental error (*σ*_12_ = *σ*_11_). We increased the standard deviation of the contaminating errors (*σ*_12_) to model outliers. REIV had the best performance (Fig. 3(b); Zamar criterion) of all the methods in the presence of outliers. These simulations suggest that errors-in-variables methods, especially REIV, perform best when normalizing transient transfection data.

**Figure 3.**
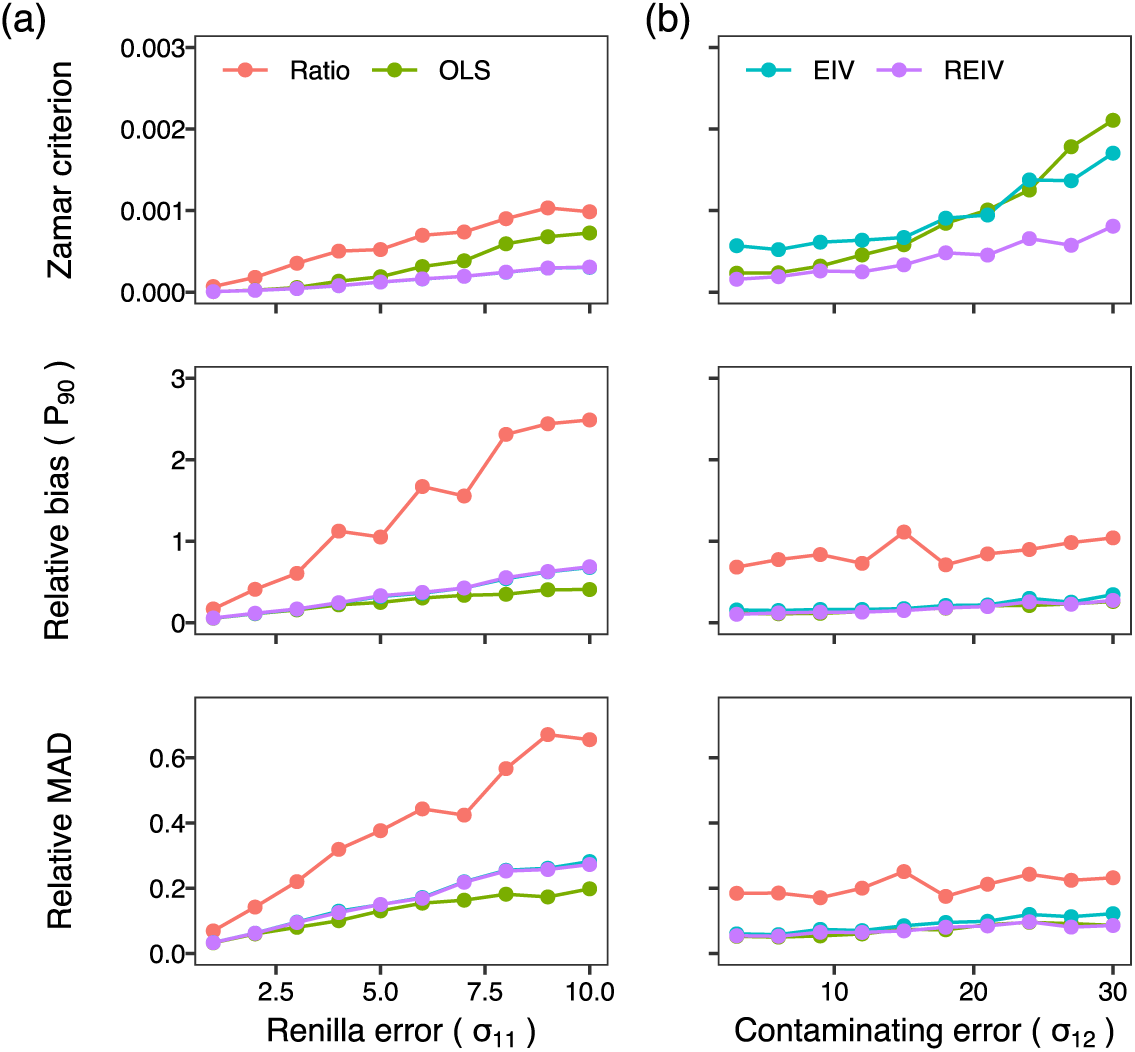
Tests of normalization methods on simulated data. The top panels plot the Zamar criterion, which is 0 for perfect inference of the true activity. The middle panels plot the 90th percentile (*P*_90_) of the relative bias. The bottom panels plot the relative median absolute deviation, a measure of precision. (**a**) Performance of the methods plotted against standard deviation of the error in *Renilla* luminescence. Mean transfection efficiency: 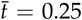. True activity: *A* = 10. Sample size: *N* = 10. There were no outliers: *σ*_12_ = *σ*_11_. Firefly errors were scaled with activity: *σ*_21_ = *σ*_22_ = 10*σ*_11_. (**b**). Performance plotted against increasing severity of outliers. The standard deviation of the contaminating *Renilla* errors *σ*_12_ was varied to simulate outliers. True activity: *A* = 3. *Renilla* errors: *σ*_11_ = 3. Firefly errors: *σ*_21_ = 9. The standard deviation of firefly contaminating errors was scaled with activity: *σ*_22_ = 3*σ*_12_. The other parameters are the same as in panel (**a**). The Zamar criterion of the ratiometric method was greater than the upper limit of the *y*-axis.

Having investigated the performance of the four methods on simulated data, we sought to test them on empirical data. We utilized a dataset of 85 measurements of firefly luminescence driven by the *Cebpa* promoter paired to *Renilla* measurements driven by the CMV promoter (Fig. 4(a)). The Luciferase assays were performed in PUER cells (Section 3.1), which have a transfection efficiency of ∼30%. We sampled the *Cebpa* promoter luminescence dataset 1000 times with replacement at sample sizes *N* = 3, 6, 10, 20, 30. Figure 4(b) shows boxplots of the normalized activity. The three regression based methods converge to a median activity of ∼13 as sample size is increased. Mirroring the results of the simulations, the ratiometric method estimates activity as ∼16 and this bias is not ameliorated by increasing the sample size. Suspecting that this bias arises from low-luminescence measurements, we sampled a subset of the data with *Renilla* luminescence greater than 8 relative luminescence units (RLU). The activity inferred by the regression based methods was unchanged as compared to the larger dataset but those inferred by the ratiometric method now agreed with the other three methods (Fig. 4(c)). This result implies that while ratiometric estimation can be biased by low-luminescence measurements, the regression based methods are robust to such data.

**Figure 4.**
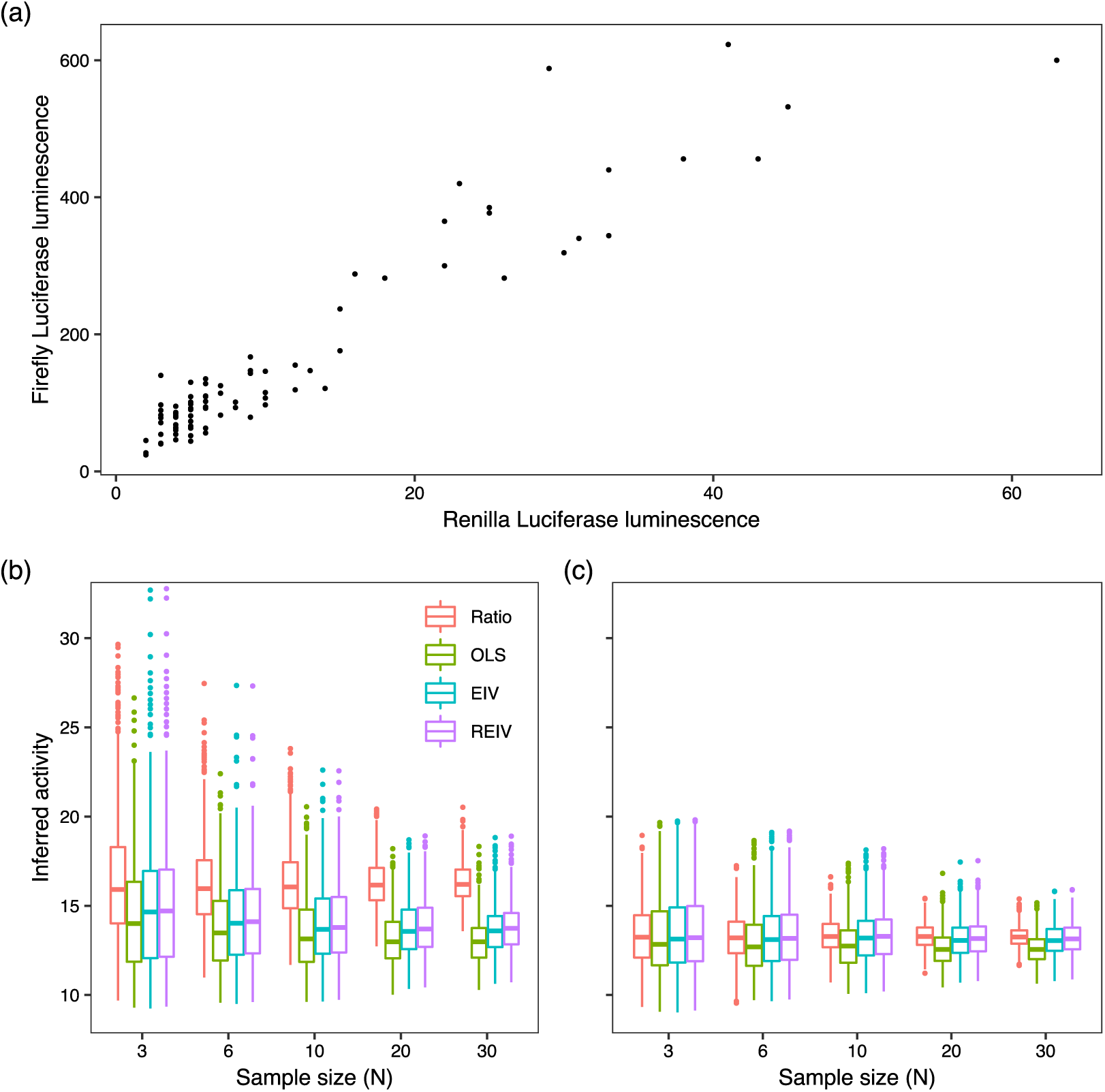
Tests of normalization methods on empirical data. (**a**) Dataset of firefly luminescence driven by the *Cebpa* promoter and *Renilla* luminescence driven by the CMV promoter in PUER cells. *N* = 85. The dataset was sampled 1000 times with varying sample sizes. Promoter activity was estimated with each method from the same exact data. (**b**) Boxplots of the inferred activities. The box lines are the first quartile, median, and the third quartile. The whiskers extend to the most extreme values lying within 1.5 times the interquartile range, and any datapoints outside the whiskers are shown as circles. (**c**) Datapoints having *Renilla* luminescence 8 RLU or less were excluded from the analysis.

Our results suggest that the commonly used ratiometric method performs adequately on reporter data from cells having transfection efficiency of at least 75% but provides inaccurate estimates in low transfection efficiency experiments. The regression based methods in contrast provide accurate inference irrespective of transfection efficiency. Among the regression based methods, REIV performs best in all conditions. Use of REIV regression for normalization could enable studies in hard-to-transfect cell lines that were not possible before. We have made R code to analyze reporter assay data using EIV and REIV available on GitHub (Sections 3.4.3 and 3.4.4).

## 3. Materials and Methods

### 3.1. Cell culture

We utilized PUER cells, *Spi1*^−/−^ cells expressing conditionally activable PU.1 protein, to assay reporter activity [3,13,14]. PUER cells were routinely maintained in complete Iscove’s Modified Dulbecco’s Glutamax medium (IMDM; Gibco, 12440061) supplemented with 10% FBS, 50*µ*M *β*-mercaptoethanol, 5ng/ml IL3 (Peprotech, 213-13).

### 3.2. Construction of the Luciferase reporter plasmid

The *Cebpa* promoter [4,15] was cloned into a pGL4.10*luc2* Luciferase reporter vector (Promega, E6651) between the XhoI and HindIII sites. The promoter was amplified from genomic DNA of C57BL/6J mice with primers TGG CCT AAC TGG CCG GTA CCT GAG CTC GCT AGC CTC GAG AAC TCC TAC CCA CAG CCG CG (Fwd) and TCC ATG GTG GCT TTA CCA ACA GTA CCG GAT TGC CAA GCT TCA GCT TCG GGT CGC GAA TG (Rev), which include 40bp of sequence homologous to pGL4.10*luc2*. PCR amplification was carried out using Q5 High-Fidelity 2X Master Mix (NEB, M0492L) following the manufacturer’s instructions. The following PCR cycling conditions were used: initial denaturation of 30s at 98C, 30 cycles of 30s at 98C, 30s at 60C, and 60s at 72C, and a final extension for 10 minutes at 72C. Gibson Assembly (GA) reactions [18] were carried out using 0.06pmol of digested vector and 0.18pmol of insert, for 60 minutes at 50C. NEB high-efficiency competent cells (NEB, E5510S) were transformed according to manufacturer’s instructions.

### 3.3. Transfection and Luciferase assays

PUER cells were transfected with *Cebpa* promoter reporter vector and *Renilla* control vector pRL-CMV (Promega, E2261) in a 1:200 ratio using a 4D-Nucleofector (Lonza). Cells were transfected with 2.26*µ*g total plasmid DNA in SF buffer (Lonza, V4SC-2096), using program CM134 and incubated for 24 hours prior to luminescence measurement. After incubation, firefly and *Renilla* luminescence were measured using the Dual-Glo Luciferase activity kit (Promega, E2920) and the DTX 880 Multimode Detector (Beckman Coulter) according to manufacturer’s instructions. Transfections were performed in at least 10 replicates.

### 3.4. Tested normalization methods

#### 3.4.1. Ratio

For each measurement, the ratio of firefly and *Renilla* luminescence was computed and averaged over the samples (Eq. 1).

#### 3.4.2. Ordinary least-squares regression

The normalized activity of the construct was determined as the slope of the line *F* = *AR*, where *F* and *R* are firefly and *Renilla* luminescence respectively, using ordinary least-squares (OLS) regression. The regression was performed using the `lm` function of `R`.

#### 3.4.3. Errors-in-variable regression

The normalized activity of the construct was determined as the slope of the line *F* = *AR* using errors-in-variables (EIV) regression [16]. EIV regression is implemented as the function eiv in the code on GitHub (https://github.com/mlekkha/LUCNORM).

#### 3.4.4. Robust errors-in-variable regression

The normalized activity of the construct was determined as the slope of the line *F* = *AR* using robust errors-in-variable regression (REIV) [17]. Since the REIV algorithm is not available in any `R` package, we implemented it in `R` as follows.

*A* was estimated by minimizing the loss function

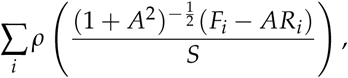

where, *R*_*i*_ and *F*_*i*_ are individual replicates of *Renilla* and firefly luminescence measurements, 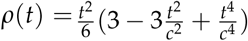 is Tukey’s loss function with *c* = 4.7, and *S* is an estimate of the scale of the residuals. The argument of Tukey’s function is the orthogonal distance of the point (*R*_*i*_, *F*_*i*_) from the regression line. Tukey’s function is bounded for large values of *t*, which limits the contribution of outliers to the loss function and ensures that the slope estimate is robust to outliers. The value of *S* was estimated by solving the equation

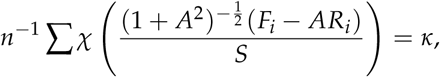

where κ = 0.05 and χ(*t*) is Tukey’s loss function with *c* = 1.56. The minimization problems were solved by the sequential least-squares quadratic programming (SLSQP) algorithm of the `NLOPTR` package of `R`, with parameters `xtol_rel` and `maxeval` set to 10^−7^ and 1000 respectively.

95% confidence intervals were estimated by bootstrapping using the `R` package `BOOT`. 999 replicates were subsampled using the ordinary simulation and the function `boot.ci` was used determine confidence intervals using the basic bootstrap method. REIV regression is implemented as the function `robusteiv` in the code on GitHub (https://github.com/mlekkha/LUCNORM).

### 3.5. Generation of simulated data

We generated simulated luminescence data that take into account sample-to-sample variation in transfection efficiency, random errors in both firefly and *Renilla* luminescence from other sources, and potential outliers. For each simulated experiment, we generated *i* = 1, …, *N* samples. The transfection efficiency of each sample was drawn from the Beta distribution,

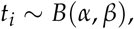

so that it varied between 0 and 1. The mean transfection efficiency of an experiment is 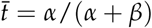. We simulated experiments with different mean transfection efficiencies (Fig. 2) by varying *α* and *β* (Table 1).

**Table 1.**
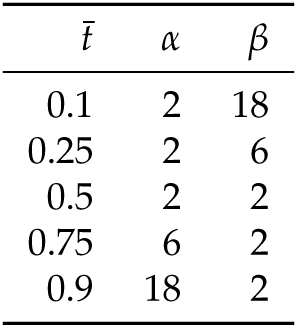
Values of Beta distribution parameters, *α* and *β*, for simulating different mean transfection efficiencies 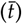.

| $\bar{t}$ | $\alpha$ | $\beta$ |
| --- | --- | --- |
| 0.1 | 2 | 18 |
| 0.25 | 2 | 6 |
| 0.5 | 2 | 2 |
| 0.75 | 6 | 2 |
| 0.9 | 18 | 2 |

The *Renilla* luminescence was computed by multiplying *t*_*i*_ with the maximal *Renilla* luminescence *R* and introducing errors drawn from a Gaussian mixture model

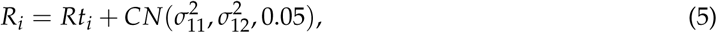

where 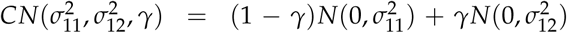 are random errors with variance 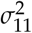 contaminated with errors with higher variance 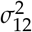. The Gaussian mixture model simulates outliers occurring with a frequency of 5%.

Firefly luminescence is computed similarly,

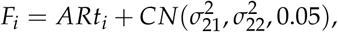

with the maximal firefly luminescence given by the product of the “true” relative activity *A* and *R*. In the simulations presented here, we assumed that the standard deviations of firefly luminescence scale with the activity so that *σ*_21_ = *Aσ*_11_ and *σ*_22_ = *Aσ*_12_.

## Author Contributions

conceptualization, A.R. and M.; software, M.; investigation, A.R. and M.; writing–original draft preparation, M.; writing–review and editing, A.R. and M.; supervision, M.; funding acquisition, M.

## Acknowledgments

PUER cells were a gift of R. Dahl.

## Abbreviations

The following abbreviations are used in this manuscript:

PUER: PU.1 Estrogen Receptor
OLS: Ordinary least-squares
EIV: Errors-in-variables
REIV: Robust errors-in-variables

## References

1. Arnold, C.D.; Gerlach, D.; Stelzer, C.; Boryn, L.M.; Rath, M.; Stark, A. Genome-Wide Quantitative Enhancer Activity Maps Identified by STARR-seq. Science 2013. doi:10.1126/science.1232542.

2. Whyte, W.A.; Orlando, D.A.; Hnisz, D.; Abraham, B.J.; Lin, C.Y.; Kagey, M.H.; Rahl, P.B.; Lee, T.I.; Young, R.A. Master transcription factors and mediator establish super-enhancers at key cell identity genes. Cell 2013, 153, 307–19. doi:10.1016/j.cell.2013.03.035.

3. Laslo, P.; Spooner, C.J.; Warmflash, A.; Lancki, D.W.; Lee, H.J.; Sciammas, R.; Gantner, B.N.; Dinner, A.R.; Singh, H. Multilineage transcriptional priming and determination of alternate hematopoietic cell fates. Cell 2006, 126, 755–66. doi:10.1016/j.cell.2006.06.052.

4. Repele, A.; Krueger, S.; Bhattacharyya, T.; Tuineau, M.Y.; Manu. The regulatory control of Cebpa enhancers and silencers in the myeloid and red-blood cell lineages. PLoS One 2019, 14, e0217580. doi:10.1371/journal.pone.0217580.

5. Stratowa, C.; Himmler, A.; Czernilofsky, A.P. Use of a luciferase reporter system for characterizing G-protein-linked receptors. Current Opinion in Biotechnology 1995, 6, 574–581. doi:https://doi.org/10.1016/0958-1669(95)80095-6.

6. Savkur, R.S.; Bramlett, K.S.; Stayrook, K.R.; Nagpal, S.; Burris, T.P. Coactivation of the human vitamin D receptor by the peroxisome proliferator-activated receptor gamma coactivator-1 alpha. Mol Pharmacol 2005, 68, 511–7. doi:10.1124/mol.105.012708.

7. Kato, M.; Sanada, M.; Kato, I.; Sato, Y.; Takita, J.; Takeuchi, K.; Niwa, A.; Chen, Y.; Nakazaki, K.; Nomoto, J.; Asakura, Y.; Muto, S.; Tamura, A.; Iio, M.; Akatsuka, Y.; Hayashi, Y.; Mori, H.; Igarashi, T.; Kurokawa, M.; Chiba, S.; Mori, S.; Ishikawa, Y.; Okamoto, K.; Tobinai, K.; Nakagama, H.; Nakahata, T.; Yoshino, T.; Kobayashi, Y.; Ogawa, S. Frequent inactivation of A20 in B-cell lymphomas. Nature 2009, 459, 712–6. doi:10.1038/nature07969.

8. Jacobs, J.L.; Dinman, J.D. Systematic analysis of bicistronic reporter assay data. Nucleic Acids Res 2004, 32, e160. doi:10.1093/nar/gnh157.

9. Fan, F.; Wood, K.V. Bioluminescent assays for high-throughput screening. Assay Drug Dev Technol 2007, 5, 127–36. doi:10.1089/adt.2006.053.

10. Minkovsky, A.; Sahakyan, A.; Bonora, G.; Damoiseaux, R.; Dimitrova, E.; Rubbi, L.; Pellegrini, M.; Radu, C.G.; Plath, K. A high-throughput screen of inactive X chromosome reactivation identifies the enhancement of DNA demethylation by 5-aza-2’-dC upon inhibition of ribonucleotide reductase. Epigenetics Chromatin 2015, 8, 42. doi:10.1186/s13072-015-0034-4.

11. Smale, S.T. Luciferase assay. Cold Spring Harb Protoc 2010, 2010, pdb.prot5421. doi:10.1101/pdb.prot5421.

12. Figueiredo, M.S.; Brownlee, G.G. cis-acting elements and transcription factors involved in the promoter activity of the human factor VIII gene. J Biol Chem 1995, 270, 11828–38. doi:10.1074/jbc.270.20.11828.

13. Walsh, J.C.; DeKoter, R.P.; Lee, H.J.; Smith, E.D.; Lancki, D.W.; Gurish, M.F.; Friend, D.S.; Stevens, R.L.; Anastasi, J.; Singh, H. Cooperative and antagonistic interplay between PU.1 and GATA-2 in the specification of myeloid cell fates. Immunity 2002, 17, 665–76.

14. Dahl, R.; Walsh, J.C.; Lancki, D.; Laslo, P.; Iyer, S.R.; Singh, H.; Simon, M.C. Regulation of macrophage and neutrophil cell fates by the PU.1:C/EBPalpha ratio and granulocyte colony-stimulating factor. Nat Immunol 2003, 4, 1029–36. doi:10.1038/ni973.

15. Bertolino, E.; Reinitz, J.; Manu. The analysis of novel distal Cebpa enhancers and silencers using a transcriptional model reveals the complex regulatory logic of hematopoietic lineage specification. Dev Biol 2016, 413, 128–44. doi:10.1016/j.ydbio.2016.02.030.

16. Casella, G.; Berger, R.L. Statistical Inference, 2nd ed.; Duxbury Press, 2001.

17. Zamar, R.H. Robust Estimation in the Errors-in-Variables Model. Biometrika 1989, 76, 149–160. doi:10.2307/2336379.

18. Gibson, D.G.; Young, L.; Chuang, R.Y.; Venter, J.C.; Hutchison, 3rd, C.A.; Smith, H.O. Enzymatic assembly of DNA molecules up to several hundred kilobases. Nat Methods 2009, 6, 343–5. doi:10.1038/nmeth.1318.

